# Unpacking fitness differences between two invaders in a multispecies context

**DOI:** 10.1101/2024.11.07.622440

**Authors:** Tomas Freire, Sten Madec, Erida Gjini

## Abstract

Ecosystems are constantly exposed to newcoming strains or species. Which newcomer will be able to invade a resident multi-species community depends on the invader’s relative fitness. Classical fitness differences between two growing strains are measured using the exponential model. Here we complement this approach, developing a more explicit framework to quantify fitness differences between two co-invading strains, based on the replicator equation. By assuming that the resident species’ frequencies remain constant during the initial phase of invasion, we are able to determine the invasion fitness differential between the two strains, which drives growth rate differences post-invasion. We then apply our approach to a critical current global problem: invasion of the gut microbiota by antibiotic-resistant strains of the pathobiont *Escherichia coli*, using previously-published data. Our results underscore the context-dependent nature of fitness and demonstrate how species frequencies in a host environment can explicitly modulate the selection coefficient between two strains. This mechanistic framework can be augmented with machine-learning algorithms and multi-objective optimization to predict relative fitness in new environments, to steer selection, and design strategies to lower resistance levels in microbiomes.

## 1 Introduction

Why do we measure fitness effects of mutations? One of the key goals is to link them to underlying evolutionary processes and the particular dynamics of alleles under selection, but also to overarching ecological and environmental processes. A typical way to quantify this is by the selection coefficient *s*. When considering an asexual (haploid) population consisting of two genotypes, a mutant and a wild-type, with respective population sizes (or densities) *N*_1_ and *N*_2_, and frequencies *p* and (1 − *p*), under the assumptions of continuous growth and no age structure, we can define the selection coefficient as 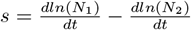, [1], which has units of *time*^−1^. The mutant will increase in frequency if *s >* 0 and decrease if *s <* 0, at a speed determined by *s*. Since the ratio of allelic frequencies is equal to the ratio of population sizes (or densities) of each genotype, we may also write 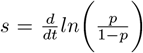. In particular, if selection is density-independent and the two genotypes do not interact, then *s* = *r*_*m*_ − *r*_*w*_ where *r* is the Malthusian parameter [1] or intrinsic rate of increase of each genotype (m, mutant; w, wild-type). In practice, *r* is estimated as the regression slope of log-population size against time in the exponential phase, i.e. at low population density (assuming no Allee effects). Such methodology is widely studied [2] and classically used in biological studies of bacterial evolution [3, 4], models to describe invader dynamics over short periods of time [5, 6], models to compare fitness of different strains [7], with applications ranging from microbial competition studies and experimental evolution, to epidemiological modeling, including assessments of viral variants during the COVID-19 pandemic [8–10].

Yet, understanding fitness differences between two entities within a complex environment, e.g. an ecosystem, presents more challenges. The cost or selective advantage of a certain mutation is strongly influenced by the environment in which the organism grows, both in its abiotic (for example, nutrient availability) and biotic (interactions with other organisms) dimensions. How organism physiology and bio-procesess in dealing with these components give rise to phenotypes, and how these phenotypes map to fitness, is a central question in evolutionary biology and ecology, with heterogeneity in selective advantage across environments a focus of modern investigations.

A key scenario of this kind arises when two invaders co-invade simultaneously an ecosystem composed of other species, for example an antibiotic-resistant and antibiotic-susceptible bacterial strain enter a multispecies microbiota context [11]. In general co-invasion, where multiple invading species or strains enter a system simultaneously, is a widespread phenomenon with significant implications in microbiology, epidemiology and agriculture. For instance, co-invasion is commonly observed in agricultural crops and plant communities [12–14], and has been important for understanding the context-dependent nature of antibiotic resistance [11, 15] and pathogen dynamics in epidemiology [16, 17].

Although it is expected that the ability of different species or strains to invade and establish themselves in a new environment depends on subtle differences in growth rates, which can be influenced by interactions with the resident species [18–20], a framework to quantify these relationships is not yet developed. In this report, we present a theory-driven model framework [21] aimed at quantifying the fitness differences between two co-invading strains, in a multi-species context. We apply our framework to a series of co-invasion experimental data with *E*.*coli* strains in mice [11], quantifying fitness costs between drug-resistant and wild-type strains. The data originate from competitive fitness assays of antibiotic resistance in the mouse gut. The antibiotic-resistant strains used carried common streptomycin (StrR (*rpsLK43T*)) and rifampicin (RifR (*rpoBH526Y*)) resistance mutations, and the double-resistant strain carried both mutations (StrR RifR (*rpsLK43T rpoBH526Y*)). These mutations have been identified in many critical pathogens, such as *Mycobacterium tuberculosis* and *Salmonella spp*., and also in pathogenic and commensal *E. coli* bacteria [22–24]. The aim of this study was to examine how interspecies interactions in the natural ecosystem comprising the mammalian gut influence the costs of antibiotic resistance in vivo, specifically, the initial co-invasion dynamics of two strains in mice that had a complex microbiota.

By harnessing this dataset, as a prototype dataset to apply our method, in a *proof-of-concept* spirit, we establish a methodology to directly link heterogeneity in selective advantage or fitness cost between two invaders, to underlying host environment multi-species composition (in this case microbiota). We first detail the mechanistic approach based on linearization of the replicator model (LR-M), highlighting its strengths and limitations, and secondly we also illustrate an approach based on machine learning algorithms, namely the Random-Forest (RF-A). Both of these approaches provide tools and insights for determining how resident polymorphic communities, in this case microbiota, shape initial invasion dynamics between two invaders.

More generally, our research aims to improve the understanding of how the multi-species composition of a host environment influences selective advantage of invaders, and whether it can be used to predict and steer competition in future environments, including for ecosystem control or biomedical applications.

## 2 Methods

### 2.1 Co-invasion in a multispecies context

The classical measurement of the selection coefficient for two strains, in a co-invasion scenario, can yield different values depending on the environment. Variations in resources, environmental conditions, and biotic or abiotic factors can all influence the outcome, as each of the two invaders may differ in relating to such gradient. A particular case of environmental variation is a multispecies context, studied, for example, by [11], where variation in co-invasion outcomes can be seen as a final consequence of host microbiota composition. Although this study reported different fitness costs for the same antibiotic resistant strain relative to the WT strain in different mice and proposed a resource-consumer model, it did not fully close the explanatory loop to link the specific microbiota variation to that in selection coefficients. Here, we aim to close this gap, using and highlighting a replicator framework [21, 25].

### 2.2 Empirical invasion of two invaders

When two strains are mixed in equal proportions (1:1) and grow independently, their relative abundances initially follow the equation:

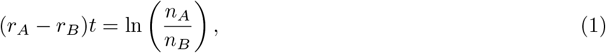

where *r*_*A*_ and *r*_*B*_ are their respective growth rates. From this linear relationship with time, we can obtain the selection coefficient between the two strains, *s* = *r*_*A*_ − *r*_*B*_.

### 2.3 A replicator equation model for multispecies dynamics

The frequency dynamics of *N* species (*z*_*i*_) is modeled using a replicator equation framework [21, 25]:

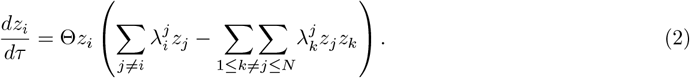

In this equation, *τ* represents time, Θ a speed constant, and 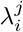 the pairwise initial growth rates between any species *i* and *j*, when *i* grows in an equilibrium set by *j* alone, also known as pairwise invasion fitness [26]. This model is particularly useful when working with microbiota data, since such data often arises from high-throughput sequencing techniques like 16S rRNA sequencing, which typically measures the relative abundance of microbial taxa.

The replicator model has an equivalent Lotka-Volterra model for *N* species with equal growth rates, and interaction matrix (*a*_*ij*_), where 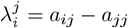 for any two species.

### 2.4 Scenario of 2 strains invading a multi-species system

The initial growth rate of an invader in a system at state **z**(*τ*) depends on both invader and system traits and is given by:

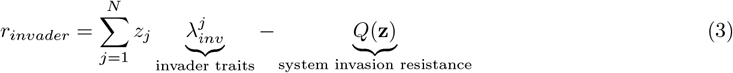

where 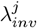, depends on invader traits, namely how this entity interacts with each resident species *j* = 1, …, *N* in the system at time of invasion. Here, *Q* depends on the system as a whole and is equal for all invaders [25]. With this approach, and under the assumption of a linear selection coefficient, we attribute this linearity to the constant frequencies of the resident species in the system during the initial invasion period.

When 2 strains, A and B, co-invade a multi-species host environment, the initial difference in their growth rates is:

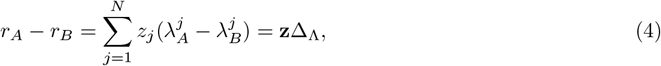

where the common quadratic term *Q* has canceled out, and where **z** is the row vector of species frequencies in the resident, and Δ_Λ_ is the column vector invasion fitness differences between the two strains relative to resident species. Assuming low initial frequencies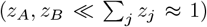, compared to those of residents, the initial influence of the two invaders can be neglected in the system. Under many realizations of such co-invasion in different resident systems, we have: **s** = *F* Δ_Λ_ where **s** = (*r*_*A*_ − *r*_*B*_)_*i*=1‥*M*_ are observed growth rate differences between A and B in *M* different hosts, and *F* the *M* × *N* matrix of their multispecies frequency compositions.

## 3 Results

### 3.1 Growth rate differences linked to microbiota compositions via the replicator

We assume the frequencies of *N* resident species are measured just prior to invasion, in *M* replicates (hosts) of the same co-invasion with 2 strains, collected in data matrix *F*. The empirical initial growth rate differential between strains *A* and *B*, is denoted by a vector **s**, which links to *F* via:

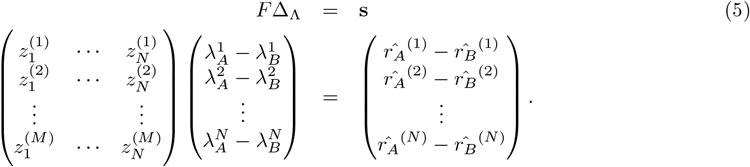

Thus, each row of *F* is the frequency vector of multispecies composition in each host (assumed constant), Δ_Λ_ the invasion fitness difference vector between the two strains, and **s** the observed selection coefficient in each host. The goal is to estimate the vector Δ_Λ_.

When *N* = *M*, we can obtain Δ_Λ_ as an exact solution of the linear system:

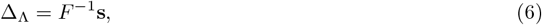

and this requires the data matrix *F* to be invertible.

When *N < M*, the number of species is less than or equal to the number of hosts, we can use least squares regression:

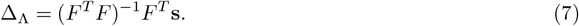

This allows us to leverage many classical statistical results regarding the uncertainty of estimation and prediction.

When *N > M*, the number of species exceeds that of hosts, and one can proceed with some suitable aggregation of species to a lower number, to reduce the number of variables, and then apply the same method.

### 3.2 From model to data: *E*.*coli* strains invade different microbiota

We extracted and prepared data from [11], and applied the model to 3 co-invasion experiments between antibiotic-resistant mutants and wild-type *E*.*coli* : 1) Rif. resistant vs. WT in single-caged mice, 2) Rif. resistant vs. WT in co-housed mice, and 3) Strep. resistant vs. WT in co-housed mice (see Supplementary Dataset and Fig.1), with microbiota resolution at the phylum level (*N* = 4 species). The species were Bacteroidetes, Proteobacteria, Verrucomicrobia and Firmicutes.

**Figure 1:**
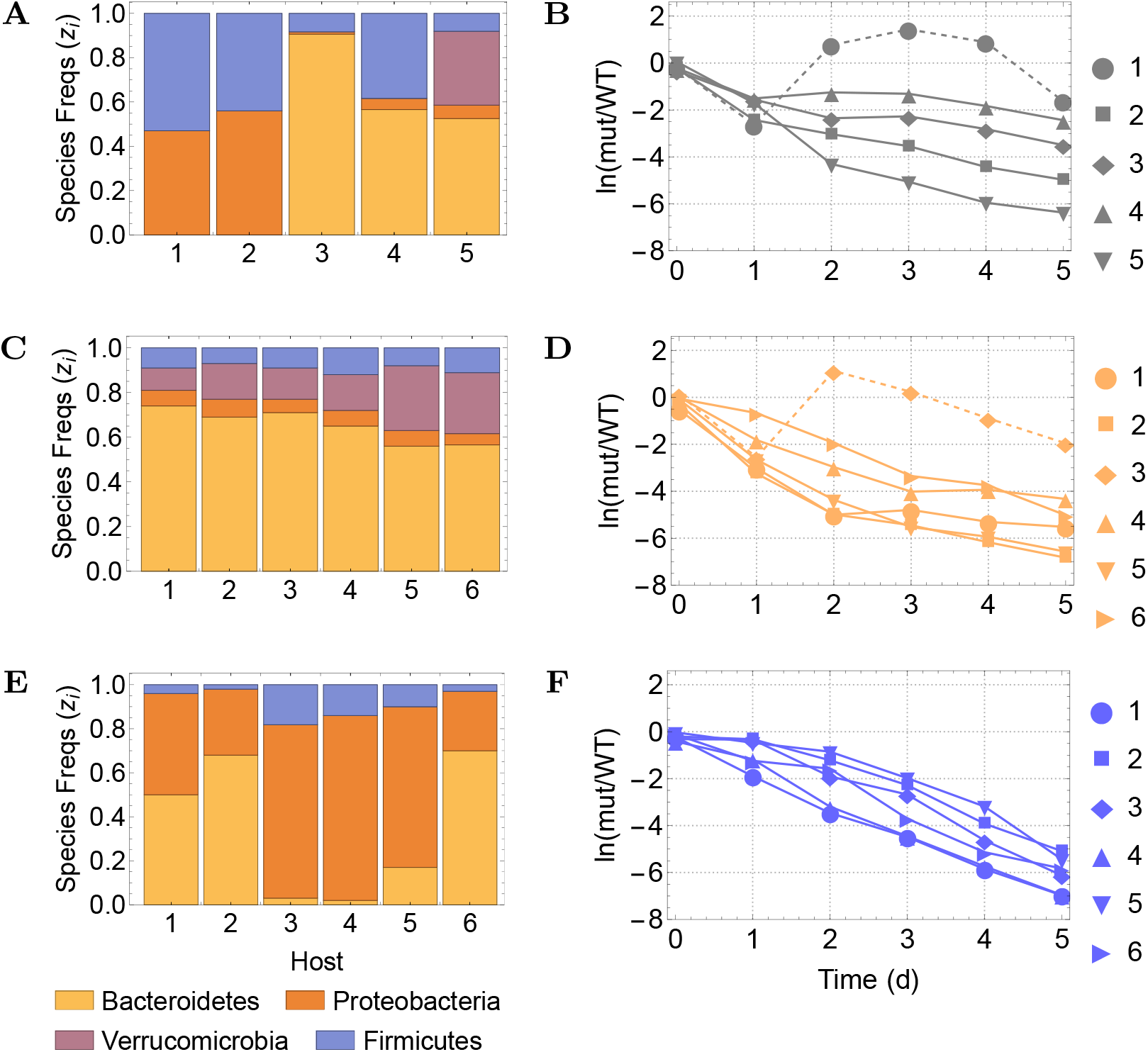
Microbiota compositions and initial relative growth between two invaders. Data publicly available and extracted from experiments with E.coli strains in mice [27]. **A-B** Data 1 (*N* = 5, *M* = 4): Mice were single-housed and injected with WT + a rifampicin-resistant mutant; **C-D** Data 2 (*N* = 6, *M* = 4): Mice were co-housed and injected with WT + a rifampicin-resistant mutant; **E-F** Data 3 (*N* = 6, *M* = 3): Mice were co-housed and injected with WT + a streptomycin-resistant mutant. Left panels show host microbiota compositions just prior to invasion. Right panels show plots of *ln*(*Res/Wt*). The slopes *r*_*A*_ − *r*_*B*_ were estimated from the linear fit with time, and their significance, assessed by an F-test. Only hosts with a significant non-zero slope were retained for analysis (mouse 1 in Data 1 and mouse 3 in Data 2 were removed, depicted in dashed lines in B,D). For details see SI dataset and SI Text 1.

We also combined data 1 and 2 under a joint regression, as they shared the same co-invaders. This led to a larger sample size (*M* = 9), hence more statistical power. Results are in Table 1, revealing the ecological interaction basis of fitness cost in these antibiotic-resistant strains. A fitness cost 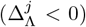 for each antibiotic-resistant strain is confirmed relative to WT, with respect to any phyla (*j* = *B, P* or *V*), except for the rifampicin-resistant strain relative to species *j* = *F*, displaying a positive signal in Data 1 and 2, separately and jointly, but without reaching individual significance.

**Table 1:** Invasion fitness differential between two invading strains of E.coli relative to host microbiota. Solution of a linear system for Data 1, and regression estimates (plus 95%CI) for Data 2 (*R*^2^ = 0.999), Data 3 (*R*^2^ = 0.999), and 1+2 jointly fitted (*R*^2^ = 0.976).

| Host microbiota<br>Species $i$ | Data 1<br>Coefficients ($\Delta_{\Lambda}^j$) | Data 2<br>Coefficients ($\Delta_{\Lambda}^j$) | Data 3<br>Coefficients ($\Delta_{\Lambda}^j$) | Data 1+2<br>Coefficients ($\Delta_{\Lambda}^j$) |
| --- | --- | --- | --- | --- |
| Bacteroidetes (B) | -0.83 | -1.78 [-10.17, 6.61] | -1.26*** [-1.5, -1.02] | -1.16*** [-1.80, -0.52] |
| Proteobacteria (P) | -2.17 | -9.17 [-73.97, 55.63] | -1.02** [-1.61, -0.44] | -2.92** [-5.08, -0.75] |
| Verrucomicrobia (V) | -2.77 | -2.00 [-8.05, 4.04] | — | -2.26* [-3.99, -0.52] |
| Firmicutes (F) | 0.22 | 8.98 [-15.92, 33.89] | -3.32* [-6.45, -0.19] | 1.07 [-1.23, 3.36] |

### 3.3 Quality of selection coefficient estimation and prediction

From the regression framework, we obtain explicitly the uncertainty around each coefficient as:

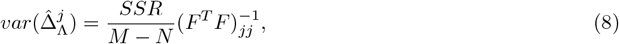

where SSR is the sum of squared residuals of the regression. Using these results, in a new microbiota composition *c*, we can predict the selection coefficient between the two strains as:

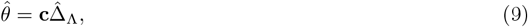

where, in mean, the standard error is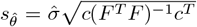, with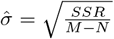, while for a particular case, 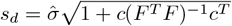, depending both on past (*F*) and current (*c*) data (See SI Text 2 for details).

### 3.4 Method validity test and potential use for microbiota engineering

#### Validity and predictive power

Having quantified Δ_Λ_, including its uncertainty, we could predict *r*_*A*_ − *r*_*B*_ when the same strains A and B invade other hosts, with any microbiota compositions. To illustrate this predictive power, we validated our linear replicator method (LR-M) on the same dataset, but through a *leave-one-out cross-validation* approach (Fig.2, SI Text 3), which confirmed good regression performance. Yet, and especially in this small dataset, sensitivity of linear regression to outliers or multicollinearity may lead to inaccurate predictions, warranting for care and preliminary tests.

**Figure 2:**
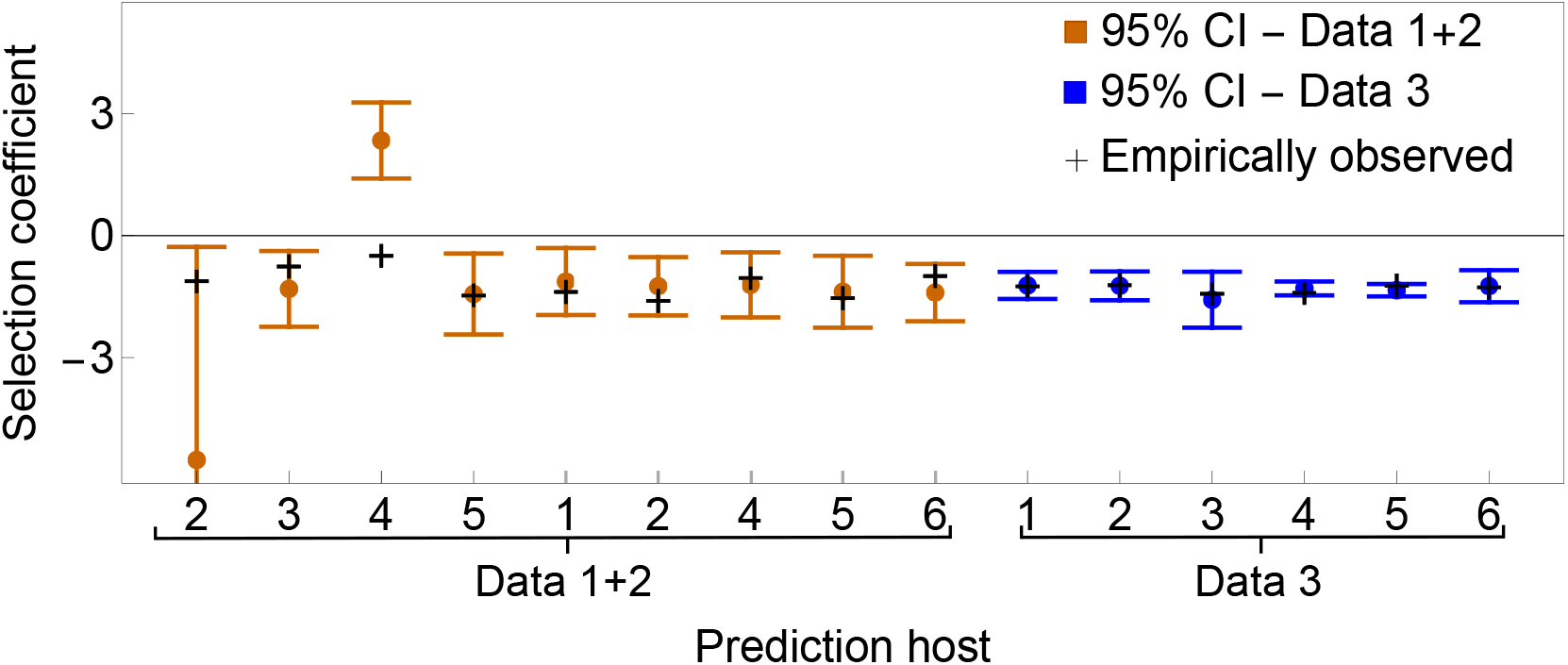
Model predictions vs. observed selection coefficients. We re-estimated invasion fitness parameters via a *leave-one-out cross-validation* procedure. For each host not used in estimation, we predicted the selection coefficient (and 95%CI), based on its microbiota composition, c. The 1 − *α* confidence interval for predicted 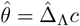 is: 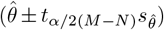 in mean, or 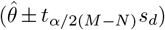 for a particular case (shown here for tests with Data 1+2 and Data 3). See also SI Text 3.

This procedure confirms that this regression-based framework predicts well selection coefficients in new hosts based on the information extracted from previous *microbiota-selection coefficients* data. Additionally, it illustrates perfectly two ways in which linear regression-based prediction can fail or be inaccurate: i) due to the presence of outliers, and ii) due to the presence of collinearity between the predictor variables (here the species frequencies). The collinearity can be captured in the 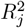 for each partial regression of a focal species on the remaining species in the dataset (see Supplementary Material, SI Text 3). For Data 1+2, when considering 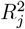 we confirmed explicitly the two cases where collinearity is very prominent: when removing mouse 2 or mouse 4 from estimation. In these two cases, in the data used for estimation, at least one variable has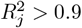, which increases the variance around the estimated regression coefficients for these species, and is likely to lead to inaccurate predictions in future cases where these species frequencies may be high.

In this particular dataset, collinearity structure is clearly the main driving factor behind inaccurate predictions for these two mice, because the other component of the variance in regression coefficients, (total variance in each variable across the sample *SST*_*j*_) is comparable between all the test cases. In contrast, for Data 3, the performance was great.

#### Microbiota engineering to steer selection

On the application front, we also explored how this method could inform the design of optimal microbiomes to steer selection coefficients towards desired regimes (Fig.3, SI Text 4). In general, determining the right microbiota compositions and diversity levels, that keep selection coefficient *s* below or above a threshold, can be framed and solved as a constrained optimization problem.

**Figure 3:**
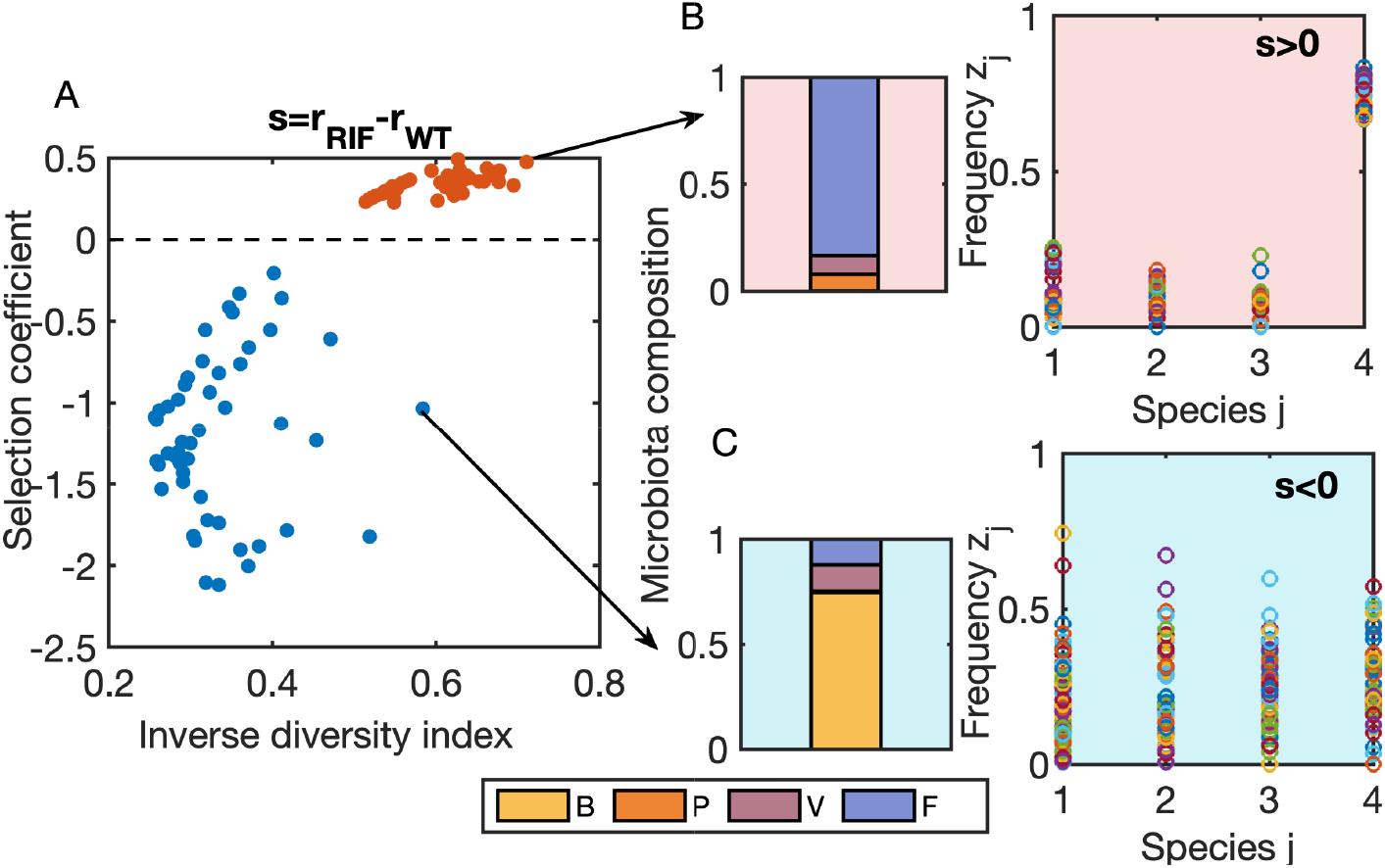
Searching microbiota compositions for target selection coefficients between invaders. We used the estimated invasion fitness differential for the Rif-WT pair of strains [27], and sought microbiota compositions yielding selection coefficients *s* in 2 target regimes: positive *s >* 0.2, and negative *s<* −0.2 (red and blue circles in **A**). We efficiently explored the space of possible solutions via minimizing 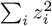 so as to favor more diversity, related to Simpson’s index. In this search, the species frequencies *z*_*i*_ potentially converge to different admissible values (**B-C**), that satisfy inequality constraints Δ_Λ_**z** *< s*_*crit*_, but vary in diversity.

In Fig. 3, we illustrate our model’s capability for use in optimization and microbiota system bioengineering. The points shown were obtained using the specific estimated coefficients of the regression for Data 1+2 (Table 1) and applying MATLAB’s fmincon function [28] to search for the microbiota compositions that satisfy the constraints of the selection coefficients being below −0.2 and above 0.2. This allowed us to explore various compositions that satisfied the constraints, starting from random initial guesses and searching the variable space in the direction of minimizing 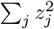. Thus, as a ‘cost’ function for the optimization search, we considered an inverse measure of diversity, which favors greater diversity related to the Simpson index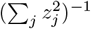, although this was not used as a strict criterion, as different solutions satisfying the constraint and varying in diversity levels were admitted for illustration.

It is important to note that the microbiota compositions illustrated do not represent the absolute optimal solution. Since this is a convex optimization problem, the true minimum could be calculated analytically. However, our use of `fmincon` was an efficient way to find points that satisfied the constraints, illustrating the wide space of potential balances between host microbiota composition and diversity, and the target selection coefficients desired. More specific optimization remains an open avenue for future investigation.

### 3.5 Alternative prediction using machine learning: the random forest algorithm

In addition to our mechanistic regression approach, we explored a model-free alternative by employing a Random Forest machine learning algorithm (RF-A) to predict the selection coefficient *r*_*A*_ − *r*_*B*_ from ‘learned’ host microbiota compositions:

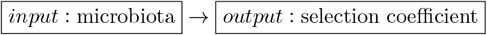

Random forests [29], one of the classic ML techniques, are an ensemble learning method used for classification and regression. They reduce variance and improve generalization by averaging predictions across multiple decision trees and have found success in the field of microbiota research [30–32]. Given the small dataset in our study, they offer robustness against overfitting and can perform well even when explanatory variables outnumber observations [33]. Unlike the proposed mechanistic model, where a linear relation between species frequencies and selection coefficients naturally arises, random forests do not assume a predefined relationship between explanatory variables (input) and the response variable (output). Instead, they infer patterns directly from the data by aggregating predictions from multiple decision trees [34]. The primary hyperparameter we control in our random forest model is the number of trees. Increasing this parameter improves stability, but also increases computational costs [35]. We implemented this algorithm in Python and used a manually specified number of trees while keeping the other hyperparameters at their default values. For more details, see SI Text 5 and SI Code.

Our first goal was to assess the model’s performance using leave-one-out cross-validation (LOO-CV), configuring the RF with 20 trees and repeating the training process 10 times to account for inherent stochasticity. Secondly, we wanted to compare the predictive performance of the replicator-based linear regression and the random forest algorithm. Our final goal was to evaluate whether different levels of microbiota resolution influence predictive accuracy in the RF setting. Results are summarized in Figure 4 and Tables 2,**??**.

**Table 2:** Relative Errors from LOOCV performance of the two approaches. To evaluate the performance of both methods, the replicator-based regression and the random-forest algorithm, we considered the relative error for each instance of prediction of true selection coefficient *s*: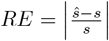, and finally, two key performance metrics by averaging over several predictions: Mean Relative Error 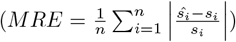 and Root Mean Squared Error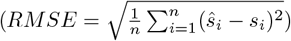. We show the absolute relative error for each point prediction using LR-M (linear replicator model) and RF-A (Random Forest algorithm) on Data 1+2 and Data 3 (rounded to two decimal places) and take their average. In gray are highlighted the cases when the point prediction by the replicator-based linear regression is closer to the empirical selection coefficient than the prediction by the Random Forest algorithm (30% of predictions for Data 1+2, and 50% of predictions for Data 3).

|  | Prediction host index | LR-M | RF-A |
| --- | --- | --- | --- |
| **Data 1+2** | 2 | 3.92 | 0.16 |
|  | 3 | 0.72 | 0.45 |
|  | 4 | 5.69 | 1.02 |
|  | 5 | 0.03 | 0.17 |
|  | 1 | 0.19 | 0.12 |
|  | 2 | 0.23 | 0.16 |
|  | 4 | 0.17 | 0.20 |
|  | 5 | 0.10 | 0.13 |
|  | 6 | 0.41 | 0.00 |
| *Global comparison* | MRE | 1.27 | 0.28 |
|  | RMSE | 2.32 | 0.41 |
| **Data 3** | 1 | 0.02 | 0.01 |
|  | 2 | 0.01 | 0.03 |
|  | 3 | 0.10 | 0.05 |
|  | 4 | 0.08 | 0.05 |
|  | 5 | 0.09 | 0.13 |
|  | 6 | 0.02 | 0.03 |
| *Global comparison* | MRE | 0.05 | 0.05 |
|  | RMSE | 0.05 | 0.06 |

**Figure 4:**
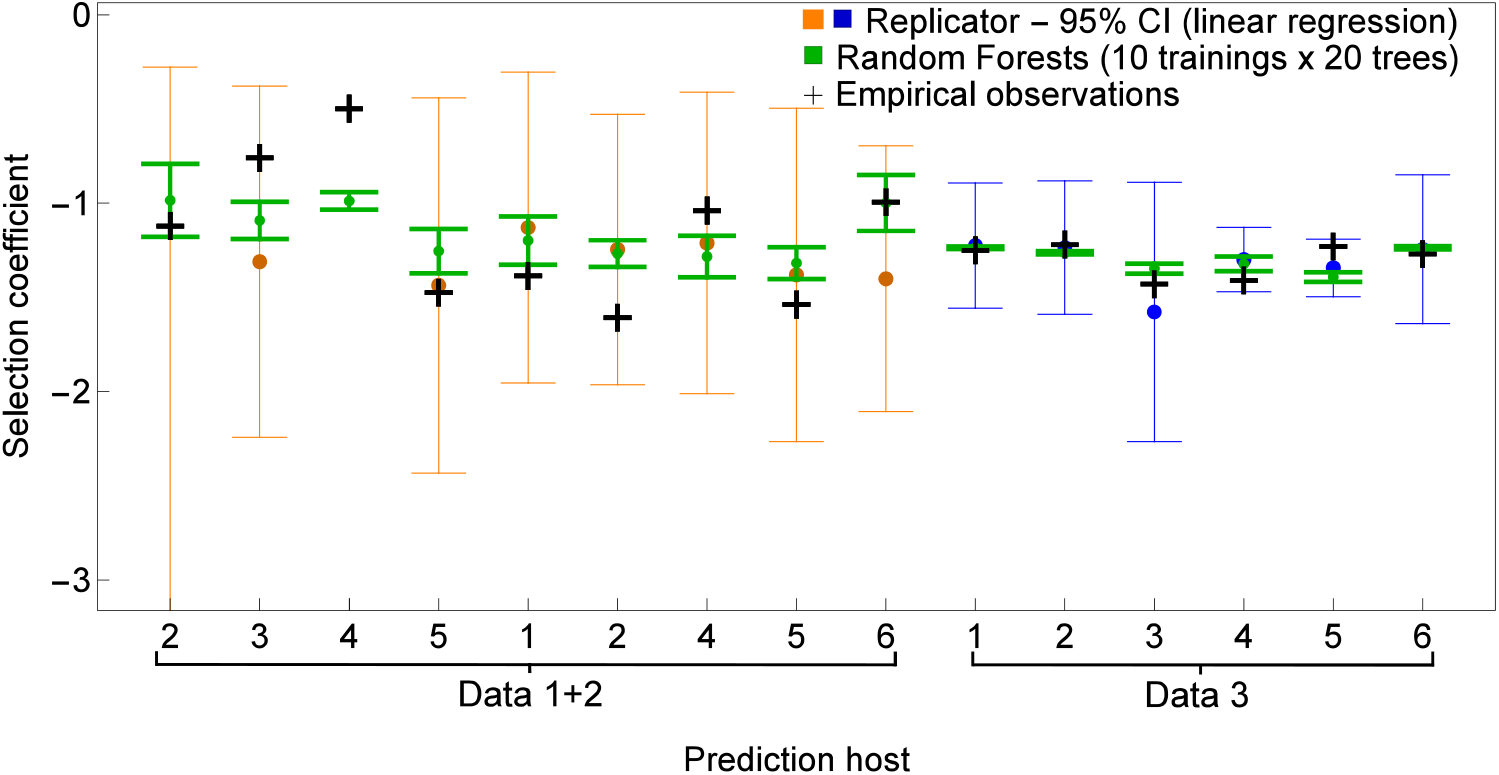
Comparing predictive performance of mechanistic vs. machine learning approach. We plot observed vs. predicted selection coefficients for the Replicator Equation-based prediction, LR-M (blue) vs. the Random Forest-based prediction RF-A (green), in the leave-one-out cross-validation approach. The actual observed selection coefficients are given by the black crosses. The figure illustrates the performance of both models, with the RF-A model achieving lower uncertainty. The interval (min, max) for the random forest algorithm was obtained by point-estimates over 10 independent trainings with 20 trees, whose mean is denoted by the green circle.

#### Comparison between the two approaches

In the Leave-One-Out Cross-Validation (LOO-CV), the model is trained on *N* − 1 data points and tested on the remaining one, iterating over all possible combinations. The RF-A algorithm was trained on the same dataset, with the microbiota frequency vector as predictors (input), and the observed selection coefficients as the response variable (output). In our case, the prediction intervals were obtained by training 10 independent models (20 trees) each time, and taking the *min* and *max* over all predicted values. For the point-estimate analysis we considered the mean of these 10 models as the point prediction from the RF-A algorithm.

We compare the quality of these predictions in Table 2. Phylum was chosen as the standard level of microbiota resolution for direct comparison with previous results, while we also explored a higher resolution level, *level 3*, though only in the RF-A case due to dimensionality constraints (Table S2).

First, when examining the results for Data 1 + 2, it is clear that the RF-A algorithm outperforms the replicator model in most predictions (Table 2, and, therefore, on average, according to both metrics. However, this trend is not observed in Data 3, where the performance metrics are nearly identical for both models. We attribute this to the machine learning algorithm’s superior ability to adapt to outliers and bypass collinearity problems of classical regression. However, when the data are of high quality, and clean of collinearity structure, such as is the case for Data 3, both approaches perform similarly well. This highlights the explanatory and predictive power of the simpler replicator framework (see SI Text 5, and Table S3 for variable significance in the RF-A setting).

However, it is worth noting that the replicator-based linear regression provides more accurate point predictions in about 30% of hosts from dataset 1+2 and 50% of hosts from dataset 3, and even in cases of lower performance, still provides confidence intervals that almost always contain the empirically-observed values of *s*, which is not the case with the uncertainty intervals obtained with RF-A. While not surprising, this data analysis seems to suggest that the linear regression framework may be more sensitive to realistic data limitations, both in terms of sample size and quality, and reflects this explicitly in the ultimate estimates. While the RF-A, machine learning algorithm, emphasizes much more strict precision, creating an impression of over-confidence, but at the expense of being sometimes less accurate (Figure 4).

Finally, when comparing the performance of the machine learning algorithm at different levels of microbiota resolution, we observe that RF-A performs better at the lower resolution level (*Phylum*) in both datasets (Table S2). This suggests that, at higher levels of resolution, the inclusion of additional species may introduce *noise* that is not relevant, and may even hamper the predictions. However, further analysis with more data is necessary before drawing definitive conclusions.

Overall, our mathematical approach clarifies and quantifies how host environment composition affects the relative competition of two invaders. With the explicit replicator model, it is possible to unpack not only the fitness cost of antibiotic resistance in terms of host microbiota species, as in this dataset [11], but more generally the fitness effects of any mutation, or any kind of invasion in a biodiverse environment.

## 4 Discussion

### Linking host microbiota to invaders’ relative fitness difference

To model invader dynamics in a multi-species ecosystem, we propose the replicator equation framework [25, 36]. It effectively and minimally describes frequency-dependent dynamics, is related to the well-known Lotka-Volterra models [37], and lends itself to the contextual nature of fitness cost between 2 strains upon co-invasion. How to connect host multi-species composition to the selection coefficient observed between two invaders? Assuming that the frequencies of resident species remain constant in the initial invasion phase, the problem simplifies to a linear system (Eqs. 5), where depending on data dimensions, either an exact solution, or optimal regression-based estimate can be found, for the vector of invasion fitness difference between the two invaders with respect to each constituent species. This method relates to and extends classical selection coefficient estimation approaches, widely used in ecology and biology, based on pure exponential growth [1, 3, 4].

We demonstrated this *proof-of-concept* on *E*.*coli* invasion data in mice with different microbiota [11], yet the framework is applicable to other multispecies composition and invasion scenarios. For example the method could be applied to *in-vivo* nasopharyngeal or lung microbiota and respiratory pathogen co-invasion [38, 39], vaginal microbiota and its protective role against viral infection [40–42], *in-vitro* invasion studies in artificially-constructed microbiomes [15], and other general multispecies environments shaping initial relative frequencies of two newcomers in the system.

We stated some obvious limitations of the method, related for example to the requirement of relatively large sample sizes (nr hosts) relative to the number of microbiota species, and those related to the sensitivity to outliers and collinearity in model variables. Yet, if data quality criteria are met, and outliers are properly handled, according to suitable statistical and information-theoretic metrics, the application of the linear regression offered by the replicator framework should pose no major problems.

As it offers an explicit quantification of the effect on each species on selection, this framework constitutes a first step in understanding underlying mechanisms, and potentially using these interactions for strategies in microbiome engineering, antibiotic resistance management, and ecosystem control.

### Mechanistic model vs. machine-learning algorithm

In recent years, machine learning techniques have gained significant traction in microbiome research, providing powerful tools to uncover complex patterns and relationships within microbial communities [43–46]. These methods offer an effective means of analyzing high-dimensional datasets often encountered in microbiota studies. Among these methods, Random Forest (RF-A), a supervised ML algorithm, has emerged as a particularly robust and flexible approach for analyzing heterogeneous microbiome data.

In this context, with the growing paradigm of AI use in research, we brought additional insights by comparing the results of our replicator-based linear regression with a classical Machine Learning algorithm, the Random Forest. Our results indicated that, depending on data quality, the RF-A may outperform the replicator model, especially in precision. However both methods perform similarly well in terms of accuracy and could be used in complementary fashion.

This dual approach enables us to compare the strength and limitations of both mechanistic models and model-free approaches in capturing the dynamics of microbial invasion. While the proposed replicator-based model with invasion-fitnesses limits the multi-species effects to be at most linearly additive for the final selection coefficient, the RF-A machine learning algorithm bypasses this constraint, and explores a wider set of possibly nonlinear effects and relationships.

So, our empirical tests indicate that if the main objective is point prediction, machine learning approaches like RF-A may be the preferred choice to connect host microbiota to invaders’ fitness differences.

While this adds to the strong predictive power of the machine learning approach, it also comes with well-known limitations, the principal one being reduced interpretability and lack of clear processes. In contrast, our linear replicator model explicitly and mechanistically quantifies how invaders interact with resident species [25], and provides confidence intervals for these estimates. This interpretability and explicit statistical foundation makes it a valuable tool for microbiota engineering, offering straightforward guidance for experimental design.

### Outlook

The next challenge lies in using the selective coefficient estimates from existing datasets to prospectively design systems that satisfy certain desired control targets. Similarly, it remains challenging to optimize diversity resolution in biotic environments for robust inference of invader parameters. There may be cases when the number of species in the microbiota composition is larger than the number of hosts, and these may require new bases of aggregation and classification, beyond genetic distance, taking into account functional diversity between members of a community. We saw that increasing the taxonomic resolution for this particular dataset in the Random Forest application did not significantly improve predictions, but of course this question deserves deeper investigation.

Finally, a key area that deserves attention is the integration of short- and long-term invasion outcomes [47], along key gradients [48], and across systems. Here, motivated also by the data [27], we treated only the initial short-term dynamics post-invasion of two invaders, and did not concern ourselves with their ultimate fate in the system. Yet, linking these initial fitness signatures to final persistence and coexistence dynamics is crucial for our understanding of long-term outcomes in ecological systems. The explicit linear approach outlined here, can be adapted to obtain the full global dynamics of the multispecies community by considering stepwise each member as an invader [25].

Furthermore, identifying the coupling parameters and processes that connect seemingly separate invasion dynamics across time and space, and environmental gradients, will provide essential unifying insights into our global quantification of fitness.

## Supporting information

SI text (integrated supplementary material)

## Acknowledgements

We thank Ermanda Dekaj for initial extraction of the data, and Isabel Gordo and Susana Vinga for helpful discussions. The work is supported by the Portuguese Foundation for Science and Technology via grant number 2022.03060.PTDC.

## Supporting Information

**SI dataset**. All microbiota composition data and selection coefficients between two invaders, extracted from [11, 27].

**SI Text**. Extended methods and references.

**SI Code**. https://github.com/tomasfreire/Unpacking_fitness_differences

