## Supplementary material for "Unpacking fitness differences between two invaders in a multispecies context": SI text (integrated supplementary material)

March 19, 2025

### 1 Notes on data preparation

We considered data reported in [1], pertaining to a series of competition experiments in mice that measured microbiota composition and invader strain levels following the invasion of both wild-type and antibiotic-resistant *E. coli* strains. From these studies, we selected three subsets of data representing three different experimental conditions, as described in [2]:

- Data 1: Rifampicin-resistant mutant vs WT in single-housed mice (5 mice);
- Data 2: Rifampicin-resistant mutant vs WT in co-housed mice (6 mice);
- Data 3: Streptomycin-resistant mutant vs WT in co-housed mice (6 mice).

#### 1.1 Microbiota Composition Analysis

The microbiota composition was analyzed at the phylum level. Initially, six phyla were detected across samples: Bacteroidetes, Proteobacteria, Verrucomicrobia, Firmicutes, Actinobacteria, and Deferribacteres. Actinobacteria and Deferribacteres were each detected in only one of the 17 mice included in the study, and at frequencies below 5%. These phyla were therefore excluded, and the remaining frequencies were normalized.

Microbiota composition was determined through 16S rRNA sequencing, specifically targeting the V4 region. The raw sequencing data in FASTQ format is available at [1] - `Fig1_Panel_F&GFinal.zip`. This data was processed using Qiime2 [3] to assess sequence quality, denoise, and classify the data. Initial quality filtering and trimming were done based on quality scores, following which denoising was performed using DADA2 [4]. Taxonomic classification of the resulting representative sequences was achieved by mapping to the Greengenes2 and SILVA databases [5, 6, 7].

The following protocol was followed for data processing and visualization:

1. Data Import and Quality Assessment: Initial data loading and quality evaluation were performed with read counts ranging from 10,000 to 45,000 per mouse.
2. Denoising with DADA2: Forward and reverse reads were truncated to 210 and 200 bases, respectively, preserving approximately 80% of the initial sequencing data.
3. Sequence Selection: A total of 148 representative sequences were identified.
4. Taxonomic Classification: representative sequences were then mapped to metagenomic data using pre-trained Greengenes2 and SILVA classifiers.

5. Results Compilation: The results were then compiled and visualized using the QIIME 2 tool, available at `https://view.qiime2.org/`.

In Figure S1, we present the Principal Component analysis (PCA) of the microbiota data, for the host with statistically significant selection coefficients. The hosts are arranged in numerical order.

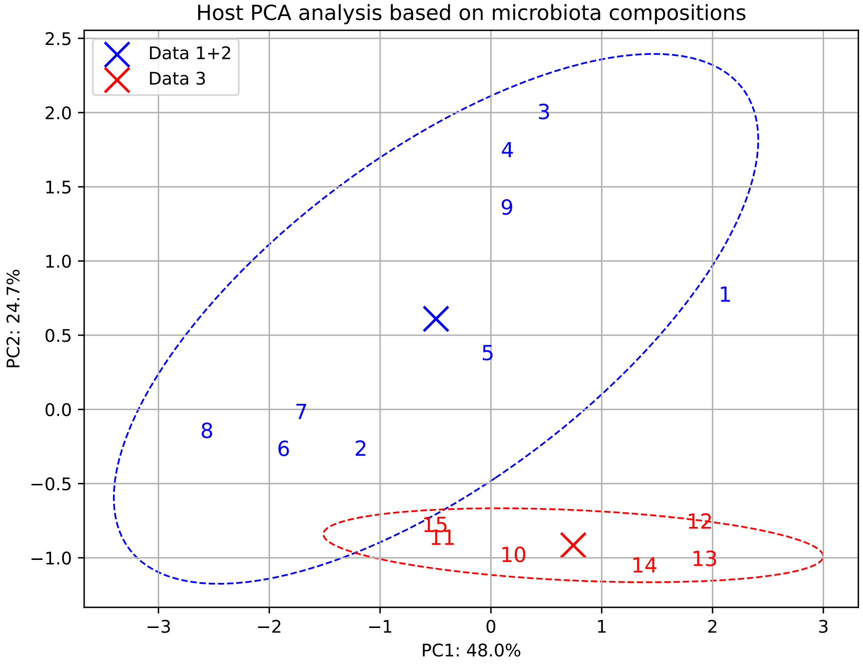


Figure S1: **Host PCA analysis based on microbiota compositions.** The microbiota information (**z**) for each host is given in detail in file *DATASI.xlsx*. Hosts in Data 1+2 were subjected to a competition (co-invasion) of a Rifampicin-resistant mutant and a Wild-type *E.coli* strain. Hosts in Data 3, were subjected to a competition (co-invasion) of a Streptomycin-resistant mutant and WT *E.coli*. Host 1 is identified as an outlier.

#### 1.2 Selection Coefficient Estimation

Selection coefficients were then estimated for each dataset by a linear regression applied to the natural logarithm of the abundance ratio between mutant and wild type, vs. time, yielding values for the slopes, under a least squares optimization approach. This data was also gathered from [1] - files `Figure1BdataFinal.xlsx` and `Figure1CdataFinal.xlsx`.

For Data 1 (Rif. mutant vs. WT in single-housed mice) and Data 2 (Rif. mutant vs. WT in co-housed mice), measurements from $t = 0$ to $t = 5$ were used, while for Data 3 (Strep. mutant vs. WT in co-housed mice), the range $t = 1 : 5$ was considered, as mutant and wild-type abundances were very similar at the two initial time points.

Hosts with slopes that were not statistically significant ($p > 0.05$) were excluded from the analysis (namely mouse 1 and mouse 3, respectively from Data 1 and Data 2), which are represented with dashed lines in Fig. 1 of the paper. All data used in the paper is given in SI dataset and the statistics of the fits are presented in table S1.

Table S1: Selection coefficients, 95% confidence intervals, and p-values for different datasets. Mouse 1 from Data 1 and Mouse 3 from Data 2 were removed due to non-significant non-zero slopes.

| Dataset | Selection Coefficient | 95% Confidence Interval | p-value | Host Index (Data 1+2) |
|---|---|---|---|---|
| **Data 1** | | | | |
| 1 | -0.03 | (-0.46, 0.40) | 0.8641 | (removed) |
| 2 | -0.88 | (-1.01, -0.75) | 0.00001 | 1 |
| 3 | -0.60 | (-0.69, -0.51) | 0.00001 | 2 |
| 4 | -0.39 | (-0.49, -0.30) | 0.0001 | 3 |
| 5 | -1.15 | (-1.36, -0.94) | 0.00003 | 4 |
| **Data 2** | | | | |
| 1 | -1.10 | (-1.38, -0.83) | 0.00015 | 5 |
| 2 | -1.27 | (-1.50, -1.04) | 0.00003 | 6 |
| 3 | -0.19 | (-0.55, 0.18) | 0.2522 | (removed) |
| 4 | -0.82 | (-0.98, -0.65) | 0.00006 | 7 |
| 5 | -1.21 | (-1.41, -1.00) | 0.00002 | 8 |
| 6 | -0.76 | (-0.92, -0.61) | 0.00005 | 9 |
| **Data 3** | | | | |
| 1 | -1.25 | (-1.40, -1.11) | 0.0001 | – |
| 2 | -1.22 | (-1.46, -0.99) | 0.00047 | – |
| 3 | -1.43 | (-1.73, -1.12) | 0.00065 | – |
| 4 | -1.41 | (-1.70, -1.13) | 0.00055 | – |
| 5 | -1.23 | (-1.83, -0.62) | 0.0076 | – |
| 6 | -1.27 | (-1.77, -0.78) | 0.0038 | – |

### 2 Notes on the regression framework and prediction intervals

#### 2.1 Statistical framework

To explain the differences in growth rates of two invading strains in a multispecies system, we considered a multiple linear regression model. In particular, in one given host $k$ (realization of the co-invasion process) we have

$$s_k = \sum_{j=1}^{N} \Delta_\lambda^j z_{k,j} + \epsilon_k, \quad k = 1, ..M \tag{1}$$

where $s_k$ is the selection coefficient observed in that host (dependent variable), $\Delta_\lambda^j$ represents the invasion fitness difference of the 2 strains, relative to species $j$ (regression coefficient), $z_{k,j}$ is the frequency of species $j$ in the host $k$ (independent predictor variable) satisfying $\sum_{j=1}^{N} z_{k,j} = 1$, and $\epsilon_k$ is one realization of the error term. In matrix form, we have

$$\mathbf{s} = F\Delta_\Lambda + \epsilon,$$

with $\epsilon \sim \mathcal{N}(0, \sigma^2)$. In this formulation, $F, \Delta_\Lambda$, and $s$, correspond to the N species frequency matrix across M different hosts ($M \times N$), the column vector invasion fitness differences between the two strains relative to resident species, and the selection coefficient values observed in each host, respectively. $F$ and $s$ come from data and we are estimating $\Delta_\Lambda$ following a least squares approach. We refer to [8] for a complete derivation of the presented results.

The least squares estimate for the coefficients of the linear regression $\Delta_\Lambda$, is given by the vector

$$\hat{\Delta_\Lambda} = (F^T F)^{-1} F^T s. \tag{2}$$

The covariance matrix for $\hat{\Delta_\Lambda}$ is given by

$$cov(\hat{\Delta_\Lambda}) = \sigma^2 (F^T F)^{-1}, \tag{3}$$

which means that the variance around each estimated regression coefficient $\Delta_\Lambda^j$ is $\sigma^2(F^TF)^{-1}_{jj}$.

An unbiased estimator for the error variance $\sigma^2$ is given by

$$\hat{\sigma}^2 = \frac{1}{M-N}\sum_{k=1}^{M}(s_k - F_k\hat{\Delta_\Lambda})^2 = \frac{SSR}{M-N}, \tag{4}$$

where $M$ is the number of observations (hosts), $N$ is the number of explanatory variables (species in each host), and $SSR$, the residual sum of squares.

**Uncertainty around estimated coefficients** The variance around the estimated coefficients can also be written as:

$$var(\hat{\Delta_\Lambda^j}) = \frac{\hat{\sigma}^2}{SST_j(1-R_j^2)} \tag{5}$$

where $SST_j = \sum_{k=1}^{M}(z_{kj} - \bar{z}_j)^2$ is the total sample variation in species $j$ frequency ($z_j$), and $R_j^2$ is the $R^2$ from regressing $z_j$ on all other species frequencies [9].

The above formula is particularly useful in highlighting how the uncertainty around each estimated coefficient increases due to three main factors: the error variance of the regression, the lack of variation in that explanatory variable (species frequency, in our case), and multicollinearity among the species frequencies.

The $1-\alpha$ confidence intervals for the estimated regression coefficient, relative to species $j$ $\Delta_\Lambda^j$ are given by

$$\hat{\Delta_\Lambda^j} \pm t_{\alpha/2,n-k}\hat{\sigma}\sqrt{(F^TF)^{-1}_{jj}}, \tag{6}$$

where $t_{\alpha/2,n-k}$ is the critical value from the Student's t-distribution with $n-k$ (in our case $n = M, k = N$) degrees of freedom. For the specific numerical example illustrated in our main text (e.g. Data 1+2), $N = 4, M = 9$ or $M = 8$ in the cross-validation part. This means for 95% CI, we use a critical value $t_{0.025,5} = 2.571$.

**Uncertainty around future observations** In a new microbiota composition $c$, the point estimate for the selection coefficient between the two strains is: $\hat{\theta} = \mathbf{c}\hat{\Delta}_\Lambda$. In mean, the standard error around this prediction is

$$s_{\hat{\theta}} = \hat{\sigma}\sqrt{c(F^TF)^{-1}c^T}, \tag{7}$$

where $\hat{\sigma} = \sqrt{\frac{SSR}{M-N}}$. However, for a particular case, the standard error is higher, because it contains also an individual realization of the random error term,

$$s_d = \hat{\sigma}\sqrt{1 + c(F^TF)^{-1}c^T}. \tag{8}$$

Thus, the $1-\alpha$ confidence interval for a new predicted selection coefficient, in a *particular* host with microbiota $c$, is given by

$$c\hat{\Delta_\Lambda} \pm t_{\alpha/2,n-k}s_d. \tag{9}$$

It is these intervals, computed for different subdivisions of the data into estimation and prediction sets, that are used in the validation procedure (Figure 2 of main text).

### 3 Methodological notes about validation procedure

To validate our results and assess the robustness of the proposed method, we used a *leave-one-out cross-validation* approach, as illustrated in Fig. 2 of the paper, using Data 1+2 and Data 3. With this approach, we systematically remove one host from the dataset, and use the remaining hosts to fit the model. The model predictive performance is then evaluated on the excluded host (predicted selection coefficient on the basis of its microbiota composition). In the case of the linear replicator model, this evaluation is done using the least squares estimate obtained with the other hosts (here 8 hosts for Data 1+2, and 5 for Data 3), (see Eq. 2). We compare how the empirically observed selection coefficient in the new host, falls within the 95% confidence interval for prediction for that particular case, computed using Eq. 9. This process is then repeated iteratively such that each host is left out once for testing.

This procedure confirms that this regression-based framework provides very good predictions in new hosts based on the information extracted from previous *microbiota-selection coefficients* data. Additionally, it illustrates perfectly two ways in which linear regression-based prediction can fail or be inaccurate: i), due to the presence of outliers, and ii) due to the presence of collinearity between the predictor variables (here the species frequencies). The collinearity can be captured in the $R_j^2$ for each partial regression of a focal species on the remaining species in the dataset. For Data 1+2 it is explicitly:

$$R_j^2 = \begin{pmatrix} \text{Removed host} & B & P & V & F \\ \text{Mouse 2} & 0.82 & \boxed{0.91} & 0.79 & 0.64 \\ \text{Mouse 3} & 0.83 & 0.76 & 0.76 & 0.83 \\ \text{Mouse 4} & 0.84 & \boxed{0.98} & 0.58 & \boxed{0.99} \\ \text{Mouse 5} & 0.70 & 0.74 & 0.57 & 0.80 \\ \text{Mouse 1} & 0.73 & 0.75 & 0.60 & 0.81 \\ \text{Mouse 2} & 0.70 & 0.75 & 0.55 & 0.81 \\ \text{Mouse 4} & 0.68 & 0.74 & 0.54 & 0.79 \\ \text{Mouse 5} & 0.68 & 0.74 & 0.53 & 0.80 \\ \text{Mouse 6} & 0.69 & 0.74 & 0.55 & 0.80 \\ \text{No host removed} & 0.71 & 0.74 & 0.58 & 0.80 \end{pmatrix}, \quad j \in \{B, P, V, F\}$$

showing two cases where it is very prominent: when removing mouse 2 or mouse 4 from estimation. In these two cases the data used for estimation suffer from high collinearity between variables (at least one variable has $R_j^2 > 0.9$, which increases the variance around the estimated regression coefficients for these species, and is likely to lead to inaccurate predictions in future cases where these species frequencies may be high. In this particular example, it is clear that this collinearity structure is the main driving factor behind inaccurate predictions for these two mice, because the other component of the variance in regression coefficients, (total variance in each variable across the sample $SST_j$) is comparable between all the test cases:

$$SST_j = \begin{pmatrix} \text{Removed host} & B & P & V & F \\ \text{Mouse 2} & 1.40 & 0.30 & 0.17 & 0.20 \\ \text{Mouse 3} & 1.03 & 0.34 & 0.17 & 0.21 \\ \text{Mouse 4} & 1.36 & 0.36 & 0.17 & 0.22 \\ \text{Mouse 5} & 1.38 & 0.36 & 0.23 & 0.20 \\ \text{Mouse 1} & 1.22 & 0.36 & 0.21 & 0.21 \\ \text{Mouse 2} & 1.26 & 0.37 & 0.23 & 0.20 \\ \text{Mouse 4} & 1.30 & 0.36 & 0.23 & 0.22 \\ \text{Mouse 5} & 1.36 & 0.36 & 0.23 & 0.20 \\ \text{Mouse 6} & 1.36 & 0.36 & 0.23 & 0.21 \\ \text{No host removed} & 1.46 & 0.40 & 0.24 & 0.23 \end{pmatrix}, \quad j \in \{B, P, V, F\}.$$

### 4 Searching microbiota compositions to satisfy given constraints

In Fig. 3 of the paper, we illustrate our model's capability for use in optimization and microbiota system bioengineering. The points shown were obtained using the specific estimated coefficients of the regression for Data 1+2 (Table 1 of the paper) and applying MATLAB's `fmincon` function [10] to search for the microbiota compositions that satisfy the constraints of the selection coefficients being below -0.2 and above 0.2. This allowed us to explore various compositions that satisfied the constraints, starting from random initial guesses and searching the variable space in the direction of minimizing $\sum_j z_j^2$.

Thus, as a 'cost' function for the optimization search, we considered an inverse measure of diversity, which favors greater diversity related to the Simpson index $(\sum_j z_j^2)^{-1}$, although this was not used as a strict criterion, as different solutions satisfying the constraint and varying in diversity levels were admitted for illustration.

It is important to note that the microbiota compositions illustrated do not represent the absolute optimal solution. Since this is a convex optimization problem, the true minimum could be calculated analytically. However, our use of `fmincon` was an efficient way to find points that satisfied the constraints, illustrating the wide space of potential balances between host microbiota composition and diversity, and the target selection coefficients desired. More specific optimization remains open.

### 5 Notes on Random Forest implementation

We employed a Random Forest regressor as provided by the `scikit-learn` Python library [11, 12]. In our experiments:

- We set the number of trees `n_estimators` to 20.
- All other parameters (e.g., `max_features`, `max_depth`) were kept at their default values.
- Training and evaluation, was done in a cross-validation process, and repeated 10 times, to mitigate the effect of random variability, inherent to the RF algorithm.

The final predictions for each data point were obtained by averaging the regressor outputs across all 10 trainings. This setup allowed us to assess the reliability of the Random Forest given a small dataset, and the average performance metrics (e.g., MRE, RMSE) provide an indication of its effectiveness.

#### 5.1 Using different taxonomic resolutions

Table S2 presents the LOOCV performance metrics (MRE and RMSE) for the RF-A at different taxonomic resolutions.

Table S2: **LOOCV Performance Metrics for different levels of resolution: Random Forest.** MRE stands for Mean Relative Error; RMSE stands for Root Mean Squared Error.

| | **Data 1+2** | | **Data 3** | |
|---|---|---|---|---|
| **Model** | **MRE** | **RMSE** | **MRE** | **RMSE** |
| Random Forest (level 2: Phylum) | 0.28 | 0.41 | 0.05 | 0.06 |
| Random Forest (level 3: more species) | 0.31 | 0.45 | 0.06 | 0.07 |

#### 5.2 Feature importance

To better compare the two approaches, we analyzed feature importance in the RF model using Mean Decrease in Impurity (MDI) [13, 14], a widely used metric for assessing variable significance in random forests. Interestingly, for Data 1+2, the variable with the largest MDI (Table S3) also had the largest confidence interval in the linear model. This might explain some performance differences between the two methods. In the linear model, this variable is unreliable due to its large confidence interval. However, in the RF model, where it is incorporated in a non-linear manner, it plays a more significant role.

Table S3: **Importance of different microbiota species in the Random-Forest approach.** We computed the MDI metric of each predictor variable for the two datasets.

| **Variable \MDI** | **Data 1+2** | **Data 3** |
|---|---|---|
| Bacteroidetes | 0.08 | 0.34 |
| Proteobacteria | 0.31 | 0.36 |
| Verrucomicrobia | 0.15 | - |
| Firmicutes | 0.46 | 0.30 |

[12] Guido Van Rossum and Fred L. Drake. *Python 3 Reference Manual*. CreateSpace, 2009.

[13] Leo Breiman. Using random forests. Technical report, University of California, Berkeley, 2002. Version 3.1.

[14] Erwan Scornet. Trees, forests, and impurity-based variable importance, 2021.
