## Supplementary material for "Unpacking fitness differences between two invaders in a multispecies context": SI data

| | **Host ID** | **Resident microbial species frequency** | | | | $r_A - r_B$ |
|---|---|---|---|---|---|---|
| | | **Bacteroidetes** | **Proteobacteria** | **Verrucomicrobia** | **Firmicutes** | |
| Data 1 | SH-R 1 | 0 | 0.47 | 0 | 0.53 | -0.03 (n.s.) |
| | SH-R 2 | 0 | 0.56 | 0 | 0.44 | -1.12 *** |
| | SH-R 3 | 0.91 | 0.01 | 0 | 0.08 | -0.76 *** |
| | SH-R 4 | 0.57 | 0.05 | 0 | 0.38 | -0.5 *** |
| | SH-R 5 | 0.53 | 0.06 | 0.33 | 0.08 | -1.48 *** |
| Data 3 | CH-R 1 | 0.74 | 0.07 | 0.1 | 0.09 | -1.39 *** |
| | CH-R 2 | 0.69 | 0.08 | 0.16 | 0.07 | -1.61 *** |
| | CH-R 3 | 0.71 | 0.06 | 0.14 | 0.09 | -0.24 (n.s.) |
| | CH-R 4 | 0.65 | 0.07 | 0.16 | 0.12 | -1.04 *** |
| | CH-R 5 | 0.56 | 0.07 | 0.29 | 0.08 | -1.54 *** |
| | CH-R 6 | 0.57 | 0.05 | 0.27 | 0.11 | -0.99 *** |
| Data 2 | CH-S 1 | 0.5 | 0.46 | 0 | 0.04 | -1.46 *** |
| | CH-S 2 | 0.68 | 0.3 | 0 | 0.02 | -0.92 *** |
| | CH-S 3 | 0.03 | 0.79 | 0 | 0.18 | -1.11 *** |
| | CH-S 4 | 0.02 | 0.84 | 0 | 0.14 | -1.43 *** |
| | CH-S 5 | 0.17 | 0.73 | 0 | 0.1 | -0.87 *** |
| | CH-S 6 | 0.7 | 0.27 | 0 | 0.03 | -1.19 *** |

Table 1: **Data.** Data used to illustrate our framework, were extracted from a previously published study [2], and describe host microbiota compositions and initial selection coefficients between two *E.coli* strains upon co-invasion. **Data 1:** SH-R mice were single-housed and injected with WT+ a rifampicin-resistant mutant. **Data 2:** CH-R mice were co-housed and injected with WT+ a rifampicin-resistant mutant. **Data 3:** CH-S mice were co-housed and injected with WT+ a streptomycin-resistant mutant. The slopes of the estimation $r_A - r_B$ from the linear fit of $ln(n_A/n_B)$ with time, are marked by $*$, $**$, or $***$, corresponding to $P < 0.05$, $P < 0.01$, and $P < 0.001$ respectively, indicating the level of significance, as assessed by a strict F-test. Only hosts with a significant non zero slope were retained in the analysis. Mouse 1 from Data 1 and Mouse 3 from Data 2 were removed. All raw data was extracted from the repository and supplementary material associated to the original study [2, 1].See also SI Text 1 for details.
